# Heat stress drives rapid viral and antiviral innate immunity activation in Hexacorallia

**DOI:** 10.1101/2024.12.19.629480

**Authors:** Ton Sharoni, Adrian Jaimes Becerra, Magda Lewandowska, Reuven Aharoni, Christian R. Voolstra, Maoz Fine, Yehu Moran

## Abstract

The cnidarian class Hexacorallia, encompassing stony corals and sea anemones, plays a critical role in marine ecosystems. Coral bleaching, the disruption of the symbiosis between stony corals and zooxanthellate algae, is driven by climate change-induced seawater warming and further exacerbated by pathogenic microbes. However, how pathogens, especially viruses, contribute to accelerated bleaching remains poorly understood. The present study utilizes the model sea anemone *Nematostella vectensis* to explore these dynamics by creating a transgenic line with a reporter gene regulated by sequences from two RIG-I-like receptor (RLR) genes involved in antiviral responses. Under heat stress, the reporter gene showed significant upregulation, indicating that these regulatory sequences are indeed responsive to thermal stress. Analyses of transcriptome data of *N. vectensis*, *Exaiptasia diaphana* (another sea anemone), and the stony coral *Stylophora pistillata* revealed stress-induced activation of a set of *bona fide* immune-related genes conserved between the three species. Population-specific differences in stress-induced transcriptional responses of immune-related genes were evident in both *Nematostella* and *Stylophora*, depending on geographic origin. In *Exaiptasia*, the presence or absence of zooxanthellae also influenced stress-induced immune gene expression. To test whether the viruses themselves may contribute to this immune response under stress, we subjected *Nematostella* polyps to variable periods of heat stress and measured the transcript levels of resident viruses as well as selected antiviral genes. While the antiviral genes responded within 1-3 hours of heat stress, viral gene expression was already upregulated within 30 minutes, suggesting that their increase might be contributing to the elevated immune response under stress, and consequentially, the further demise of organismal homeostasis. These findings highlight the complex interplay between environmental stress, viruses, immune responses, and symbiotic states in Hexacorallia. Better understanding of these mechanisms could provide insights into the role of immune pathways in coral resilience and bleaching in a changing climate.

## Introduction

Coral reefs are a habitat supporting at least 25% of all marine species (Fisher, et al. 2015; Hoegh-Guldberg, et al. 2019). They are formed by members of the order Scleractinia, also known as stony corals, which are dependent on close symbiosis with zooxanthellae, dinoflagellates in the family Symbiodiniaceae that reside within their gastrodermis (Fransolet, et al. 2012; LaJeunesse, et al. 2018; Jacobovitz, et al. 2023). Coral bleaching, driven by symbiont loss, has escalated alarmingly in recent decades, with global events increasingly frequent and severe (Sully, et al. 2019; van Woesik, et al. 2022), leading to extensive habitat loss (Reimer, et al. 2024)

The loss of these symbionts is often ultimately resulting in the death of the coral host starting as a phenomenon known as “coral bleaching”. This phenomenon has been happening in the last several decades at alarming rates with global bleaching events becoming more frequent and more extreme, resulting in the loss of complete marine habitats (Reimer, et al. 2024). The exact causes underlying coral bleaching are still under debate and this seems to be a complex phenomenon that may be attributed to numerous different factors acting in parallel (Rädecker, et al. 2021; Helgoe, et al. 2024; Schlotheuber, et al. 2024). Yet, there is a growing consensus that rise of water temperature is a major cause for this via induction of coral stress (Cziesielski, et al. 2019) that leads to the collapse of nutrient cycling between the algal symbiont and coral host and the buildup of reactive oxygen species (ROS) triggering oxidative damage (Downs, et al. 2002; Rädecker, et al. 2021; Rädecker, et al. 2023). Another potential cause frequently suggested to contributing to the phenomenon of bleaching is pathogenesis, mostly by bacteria (Rosenberg, et al. 2007). How exactly the stress and/or pathogenesis result in the loss of zooxanthellae is still being investigated and one of the major approaches in the field has been differential gene expression analysis, comparing healthy and bleached corals (DeSalvo, et al. 2008a; Seneca, et al. 2010; Bellantuono, et al. 2012; Pinzón, et al. 2015; Seneca and Palumbi 2015). Multiple studies in stressed and bleached corals observed strong upregulation of genes that are homologous to known immune-related genes from vertebrates (DeSalvo, et al. 2008b; Kenkel, et al. 2014; Rosic, et al. 2014; Pinzón, et al. 2015; Wang, et al. 2024). Thus, current evidence suggests a potential link in corals between stress, pathogenesis, and immunity. Yet, one can suggest two alternatives, non-mutually exclusive scenarios: in the first, heat stress induces pathogenesis that in turn induces the immune response. Alternatively, in the other scenario, the immune system of corals evolved to be upregulated upon heat stress as this would be advantageous against pathogens that tend to be more active under heat stress.

Although work on coral in their natural environment is invaluable to study bleaching, a detailed cell biological understanding cannot be achieved without manipulative experiments in a controlled lab environment, which prompts the use of model organisms (Weis, et al. 2008). Scleractinia are part of the cnidarian class Hexacorallia, which includes also sea anemones (Ruppert, et al. 2004). Sea anemones offer practical advantages as laboratory models due to their ease of culture and controlled reproduction. Some were developed into lab model organisms for answering questions in evolutionary developmental biology (“evo-devo”) and ecology (Darling, et al. 2005; Baumgarten, et al. 2015; Rentzsch, et al. 2017; Al-Shaer, et al. 2021; Röttinger 2021; Jacobovitz, et al. 2023; Roberty, et al. 2024). The two arguably most developed sea anemone model species are *Nematostella vectensis* and *Exaiptasia diaphana* (previously called *Aiptasia pallida*, frequently referred to as Aiptasia). While both *N. vectensis* and *E. diaphana* are separated by roughly 500 million years from stony corals (Erwin, et al. 2011; Shinzato, et al. 2011), the latter harbors zooxanthellae in their gastrodermis just like corals, making it a useful model for studying hexacorallian-dinoflagellate symbiosis (Jacobovitz, et al. 2023; Roberty, et al. 2024).

*N. vectensis* lives in distinct populations spread across isolated brackish lagoons in the Atlantic coast of North America from Canada to southern USA (Reitzel, et al. 2013). There is little gene flow between these populations and yet we have shown that despite highly similar genomic sequences (Smith, et al. 2023), they evolved different transcriptional responses to environmental stress (Sachkova, et al. 2020). Similarly, we have shown that populations of the stony coral *Stylophora pistillata* originating either from the Gulf of Aqaba (GoA) at the northern point of the Red Sea or from the central Red Sea (CRS), exhibit different transcriptional responses to heat stress (Voolstra, et al. 2021), but it is unclear whether they form distinct genetic populations (Buitrago-López, et al. 2023). Similar trends were also reported between species (Voolstra, et al. 2023). Interestingly, allelic variability was reported for the key immune- and stress-related transcription factor NF-κB between populations of *N. vectensis* (Sullivan, et al. 2009). Thus, interspecific and intraspecific genetic variation in Hexacorallia may drive different responses to environmental cues, possibly reflecting local adaptation. Yet, whether this variation has effect on immunity is currently unknown.

A recent study showed that under heat stress conditions phagocytosis activity is increased in hexacorallians (Eliachar, et al. 2022). Moreover, key players in immune response are downregulated in hexacorallians during symbiosis with zooxanthellae (Mansfield and Gilmore 2019). While much attention was given to the effect of bacteria on the hexacorallian immune response (Bourne, et al. 2009; Mao-Jones, et al. 2010; Munn 2015), far less attention was provided for viral agents. Yet, there are indications that upon heat stress, herpes-like viruses present in the coral genome are activated (Vega Thurber, et al. 2008). Moreover, there are indications that viruses that are pathogenic to the zooxanthellae become more productive under higher water temperatures (Levin, et al. 2016; Howe-Kerr, et al. 2023). Thus, viruses and the immune response that defends against them may play an underappreciated role in shaping symbiosis stability in Hexacorallia.

We have started revealing the antiviral innate immune system of *N. vectensis* by characterizing the transcriptomic response of this species to double-stranded RNA (dsRNA), a pathogen-associated molecular pattern (PAMP) typifying most viruses (Lewandowska, et al. 2021). We have shown that this viral hallmark is detected by the cytosolic receptor RIG-I-Like Receptor b (RLRb), a homolog of the mammalian RIG-I and MDA5 receptors that is conserved in all hexacorallians (Lewandowska, et al. 2021; Iwama and Moran 2023). The transcriptional response we reported is diverse and includes upregulation of homologs of genes of the vertebrate interferon response as well as of those known to participate in antiviral RNA interference (RNAi) in protostomes, such as insects and nematodes. Similarly, wide transcriptional response was detected in *N. vectensis* upon immune system activation by 2’3’-cGAMP. This cyclic dinucleotide is known to serve as a signaling molecule in the antiviral immune system of vertebrates, and this study highlighted that concomitantly homologs of antibacterial genes are upregulated by this challenge in this hexacorallian species (Margolis, et al. 2021). Currently, very little is known about how the immune systems of other hexacorallians react to viral PAMPs.

In the current work we have set to explore how stress is linked to antiviral innate immunity by analyzing existing transcriptomic resources from three major hexacorallian models, the sea anemones *E. diaphana* and *N. vectensis* and the stony coral *S. pistillata*. Additionally, we utilized transgenesis tools established for *N. vectensis* (Renfer, et al. 2010; Renfer and Technau 2017) as well as the knowledge about its core virome (Lewandowska, et al. 2020) to further explore this intriguing link.

## Materials and Methods

### Sea Anemone Culture

*N. vectensis* early life stages (embryos, larvae, and primary polyps) were grown in the dark at 22 °C and in salinity of 16 ‰ artificial seawater, whereas juveniles were grown at 18 °C. From two weeks after fertilization, the polyps were fed with *Artemia salina* nauplii three times a week. Induction of gamete spawning was performed as previously described (Genikhovich and Technau 2009) For performing microinjection, the fertilized eggs were separated from the egg package by incubation in 3 % L-cysteine (Merck Millipore, Burlington, MA). For all other usage the fertilized eggs were left for three days at 22 °C until the developed planulae was spontaneously released from the gelatinous egg sack. All *N. vectensis* individuals used in this study were of the common lab strain originating from Rhode River, Maryland, USA (Hand and Uhlinger 1992).

### Cloning and Transgenesis

The promotor regions of *NveRLRa* (scaffold_15:1,087,962-1,091,460; coordinates are taken from (Putnam, et al. 2007)) and *NveRLRb* (scaffold_40:697,555-699,446) genes were amplified by employing the Advantage 2 polymerase (Takara-Bio, Kusatsu, Japan) from a genomic DNA template. The backbone vector for these transgenesis experiments was a pCR2.1 vector (Thermo Fisher Scientific, Waltham, MA, USA) modified by the Technau lab (University of Vienna, Austria) holding a cis-regulatory element, driving a gene encoding the mCherry fluorescent reporter protein (Shaner, et al. 2004) and a SV40 viral sequence for polyadenylation signaling (Renfer, et al. 2010; Renfer and Technau 2017). The PCR fragments were amplified so they will carry *PacI* and *AscI* cloning sites which are included in the vector. After overnight digestion with the two restriction enzymes (New England Biolabs, Ipswich, MA, USA) ligation of the vector and insert was performed with T4 DNA Ligase (New England Biolabs) and the ligation products was chemically transformed to NEB5α™ Competent Cells (New England Biolabs) according to the manufacturer’s protocol. Purification of plasmids was done by HiSpeed Plasmid Midi Kit (QIAGEN N.V., Venlo, Netherlands) and validated by Sanger sequencing (performed at the Genomic Technologies Center of the Hebrew University of Jerusalem). The plasmid of each RLR gene was injected into *N. vectensis* zygotes along with the meganuclease I-*Sce*I (New England Biolabs) to enable genomic integration as previously described (Renfer, et al. 2010; Renfer and Technau 2017). Transgenic animals were visualized under an SMZ18 stereomicroscope equipped with a DS-Qi2 camera (Nikon, Tokyo, Japan) and positive animals were sorted and grown to the adult stage. To sort the positive animals, we incubate planulae at 37°C for 24 h to stimulate the gene expression of the NveRLRa and NveRLRb resulting in the expression of mCherry (see Results). In each generation, we crossed transgenic animals with Wild type animals and tested the offspring to find positive animals. Then we crossed the positive siblings until we got adult positive homozygous animals (F3). Sequences of all used primers are provided in Supporting Information table S1.

### Injection of Viral Mimics to transgenic animals

To initiate the response of the antiviral immune system in *N. vectensis*, we used dsRNA as a mimic of viral RNA. We used 6.25 ng/ml of high molecular weight (HMW) Polyinosinic:polycytidylic acid (poly(I:C)), a commercially available synthetic dsRNA, in 0.9 % NaCl (Invivogen, San Diego, CA, USA) with an average size of 1.5–8 kb, and 0.9 % NaCl as a control. The concentration was selected after a series of injections that was revealed to be the most efficient to test the antiviral immune response. Higher concentrations resulted in high mortality and aberrant zygote morphology after 24 h. In each experiment, 150–200 zygotes were injected per group and kept at 22 °C. Transgenic animals were visualized under an SMZ18 stereomicroscope equipped with a DS-Qi2 camera (Nikon).

### Dynamic heat stress assay

Wild type adult *N. vectensis* polyps were incubated for different time periods at 37 °C (basal condition 0 h, 0.5 h, 1 h, 3 h, 6 h, 24 h, 48 h). Per each biological replicate, 4-6 animals were flash-frozen in liquid nitrogen, grinded, and stored at −80 °C until further processed. We conducted the RNA extraction using Trizol (Thermo Fisher Scientific) according to the manufacturer’s protocol. cDNA was constructed using iScript cDNA Synthesis Kit (Bio-Rad, Hercules, CA, USA) according to the manufacturer’s protocol. Real-time PCR was prepared with Fast SYBR Green Master Mix (Thermo Fisher Scientific) on the QuantStudio 3 Real-Time PCR System (Thermo Fisher Scientific). The relative expression was calculated by the relative standard curve method, the relative comparison was to the basal condition. We converted the Ct value to fold change (2^^x^) and compared the ratio between the increase in the viral load or the gene expression level to the basal condition state. Sequences of all primers and statistical tests are provided in Supporting Information tables S2 and S3.

### Virome assembly

The assembly of the viromes of *N. vectensis* animals from Ft. Fisher, North Carolina and Sippewissett, Massachusetts, was based on previously published transcriptomic data (Smith, et al. 2023). The filtering, processing and assembly of data followed the established protocol we previously used for the virome of the *N. vectensis* main lab population (Lewandowska, et al. 2020). Sequences of the viruses from the *N. vectensis* viromes are provided in Supporting Information File1 and File2.

### Differential gene expression and viral load analyses

To assess the impact of heat stress on immune system activation, we reanalyzed data from five previously published studies focused on species within the phylum Cnidaria. These studies share a common emphasis on examining transcriptomic responses to natural environmental stressors, particularly heat stress. Three of the studies involve populations of the sea anemones, *N. vectensis* (Sachkova, et al. 2020; Weizman, et al. 2021) and *E. diaphana* (Cleves, et al. 2020). The remaining two studies examine the stony coral species *S. pistillata* (Savary, et al. 2021; Voolstra, et al. 2021).

### Orthology analysis to identify immune-related genes

We curated 56 *bona fide* immune-related genes significantly upregulated following poly(I:C)-induced antiviral challenge in *N. vectensis*. polyI:C is a mimic of long viral dsRNA and a primary ligand for the vertebrate RLR melanoma differentiation-associated protein 5 (MDA5) (Lewandowska, et al. 2021). In the five studies we used for our analyses, these genes were used as markers to identify the triggered transcriptomic immune response when the individuals were subjected to heat stress. Since these genes were originally identified in *N. vectensis*, an orthology analysis was conducted to identify their corresponding orthologs in the genomes of *E. diaphana* and *S. pistillata*. OrthoFinder v. 2.4.0 (Emms and Kelly 2019) which combines BLAST and phylogenetics for high accuracy was used to identify orthogroups and orthologs between the *E. diaphana* and *S. pistillata* genomes. Additionally, the identified orthologs were analyzed for their domain structures using InterProScan (https://pubmed.ncbi.nlm.nih.gov/24451626/) to validate and confirm the accuracy of the orthology inference. Only sequences that exhibited similar domain architectures across the genomes of the three species were accepted as orthologous for the corresponding immune-related genes. The list of immune-related genes for *N. vectensis* and their respective orthologs in *E. diaphana* and *S. pistillata* are provided in Supporting Information tables S4, with the corresponding sequences available in table S5.

### RNA-Seq data processing

Raw reads were downloaded from five studies and were analyzed using an identical strategy to identify immune-related genes that are up-regulated in animals subjected to heat stress compared to control animals. The quality of reads was checked with FastQC (Andrews 2010) before and after read trimming with Trimmomatic version 0.36 (Bolger, et al. 2014). Reads were aligned to the *N. vectensis* (Putnam, et al. 2007), *E. diaphana* (Baumgarten, et al. 2015) and *S. pistillata* (Voolstra, et al. 2017) genomes using STAR under default alignment parameters (Dobin, et al. 2013). Read counts matrices were generated using feature counts (Liao, et al. 2014) using default parameters. Differentially expressed genes (DEGs) were determined based on Deseq2 (Love, et al. 2014) and EdgeR (Robinson, et al. 2010) packages. Genes with log_2_ fold changes ≥ 1 and adjusted P-value ≤ 0.05 were considered to have a significant differential expression. The approach to inferring DEGs in the Weizman et al. (2021) study differed from the other four studies, as it did not include biological replicates. We identified differentially expressed genes using the methodology, implemented in the R package, NOISeq (Tarazona, et al. 2011). Essentially, NOISeq simulates technical variability to compensate for the lack of biological replicates, allowing it to estimate the probability of differential expression even when true biological variability is not directly observed. Genes with a q = Probability (differential expression) of 0.9 and a M (which is the log2-ratio of the two conditions) of 1 were considered differentially expressed.

Volcano plots were generated to visualize the results of differential expression in *N. vectensis*, while heatmaps were used to display the log2 fold change (LFC) in immune-related gene expression across baseline and stress temperatures for each of the studies analyzed. If any of the 56 evaluated immune-related genes did not meet the defined thresholds, it was assigned a value of 0 and represented as dark cyan in the heatmap.

### Viral load quantification and comparison

To better understand how heat stress affects the immune system, we compared viral loads at baseline and stress temperatures in *N. vectensis* and *E. diaphana*. For *Nematostella vectensis*, we used data from Sachkova et al., (2020). To quantify viral load, raw reads from each replicate within the Massachusetts and North Carolina populations were mapped using Bowtie2 (Langmead and Salzberg 2012) and counted using SAMtools (Li, et al. 2009). Mapping was also conducted against the complete virome of *N. vectensis*, as identified in a previous study (Lewandowska, et al. 2020). For *E. diaphana*, we utilized data from Cleves et al., 2020. The raw reads were mapped and counted following the same approach as for *N. vectensis* against the published core virome of *E. diaphana* (Bruwer and Voolstra 2018). Viral loads in *E. diaphana* were compared between baseline temperature (27 °C) and heat stress temperature (34 °C) after 3 hours. To determine whether the differences in viral loads between conditions (control and stress temperatures) for each species were statistically significant, we performed a t-test. Raw reads mapped to each virus in the respective viromes of each species were normalized to the total library size of each replicate. Statistical significance was defined as a p-value of less than 0.05. The virome sequences for *N. vectensis* and *E. diaphana* are available in Supporting Information file2 and table S6, respectively. The raw data used for the statistical tests and the corresponding figures are provided in Supporting Information table S7 for *N. vectensis* and Supporting Information table S8 for *E. diaphana*.

## Results

### *RLRa* and *RLRb* promoters are sensitive to a viral PAMP and stress

We generated transgenic *N. vectensis* lines that harbor in their genome a transgenic cassette with a gene encoding the mCherry fluorescent protein (Shaner, et al. 2004) downstream of the putative regulatory regions of the *RLRa* gene we predicted based on its histone marks (**Fig. 1 A-B**) (Schwaiger, et al. 2014). While the transgenic cassette did not drive mCherry expression in the majority of the animals under normal conditions (**Fig. 1C**), injection of the viral mimic poly(I:C) drove strong fluorescent signal within 24 hours post-injection (**Fig. 1D-E**). Similar results were obtained with a transgenic cassette containing the regulatory regions of *RLRb* (**Fig. 1 A, F-I**). The strong increase in transgene expression reflects the fact that both *RLRa* and *RLRb* are upregulated upon poly(I:C) injection (Lewandowska, et al. 2021). Unexpectedly, we noticed that some of the animals injected with NaCl (a control group for the microinjection itself) also exhibited fluorescent expression, albeit at the lower level and frequency compared to the poly(I:C) group. This led us to hypothesize that stress caused by the mechanics of the microinjection and the injury it causes promotes the expression of *RLRa* and *RLRb*. To test this notion, we exposed embryos of both transgenic lines to heat stress of 37 °C. Indeed, these embryos exhibited noticeable fluorescence when compared to the control group (**Fig. 2**). Interestingly, the fluorescence was maintained for several days after the incubation.

**Figure 1.**
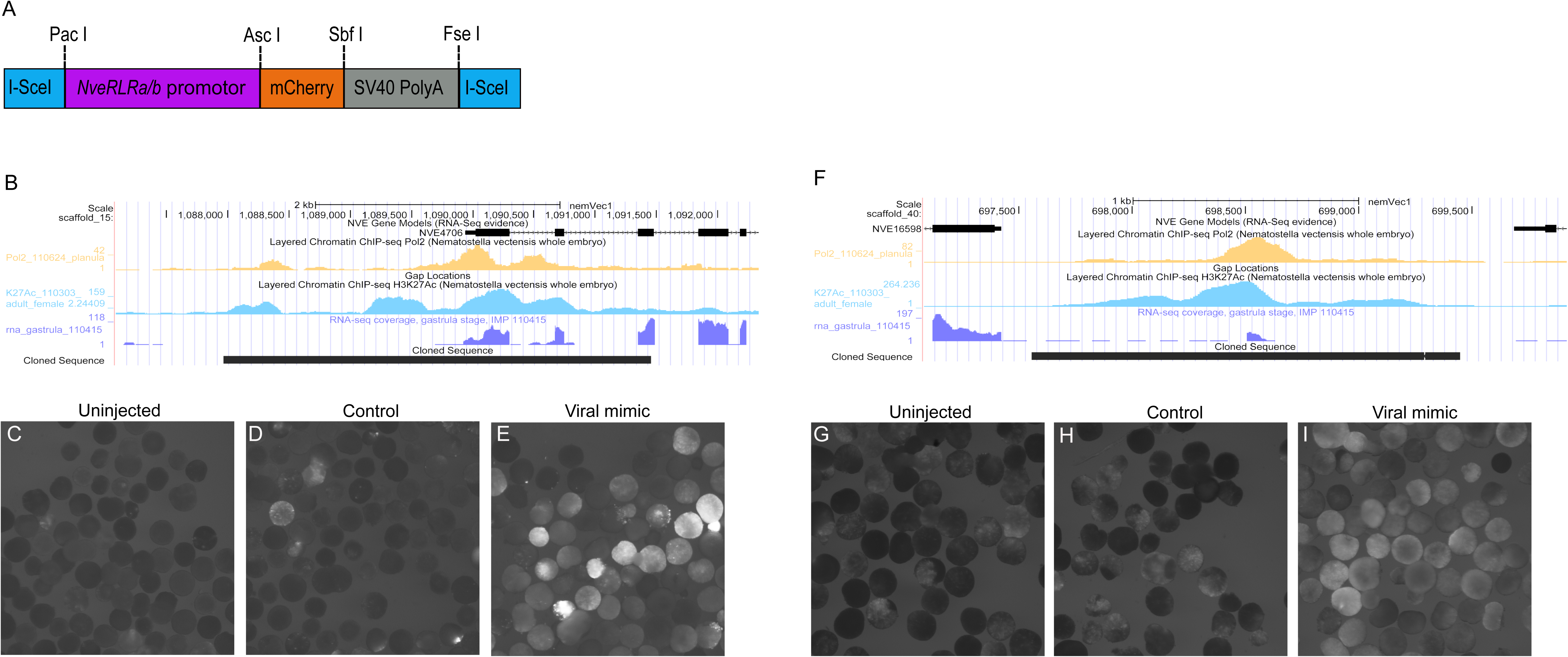
Generation of the transgenic *RLRa* and *RLRb* reporter lines and their response to dsRNA microinjection. (**A**) The design of the microinjected cassette used to generate the transgenic *N. vectensis* lines. (**B**) Features of the genomic segment containing the *RLRa* regulatory region. (**C-E**) the expression of mCherry in the *RLRa* transgenic reporter line. (**C**) uninjected animals, (**D**) NaCl injected animals, (**E**) poly(I:C) injected animals. (**F**) Features of the genomic segment containing the *RLRb* regulatory region. (**G-I**) the expression of mCherry in the *RLRb* transgenic reporter line. (**G**) uninjected animals, (**H**) NaCl injected animals, (**I**) poly(I:C) injected animals The *RLRa* transgenic reporter line was documented 19 hours after fertilization, and the *RLRb* transgenic reporter line was documented 24 hours after fertilization.

**Figure 2.**
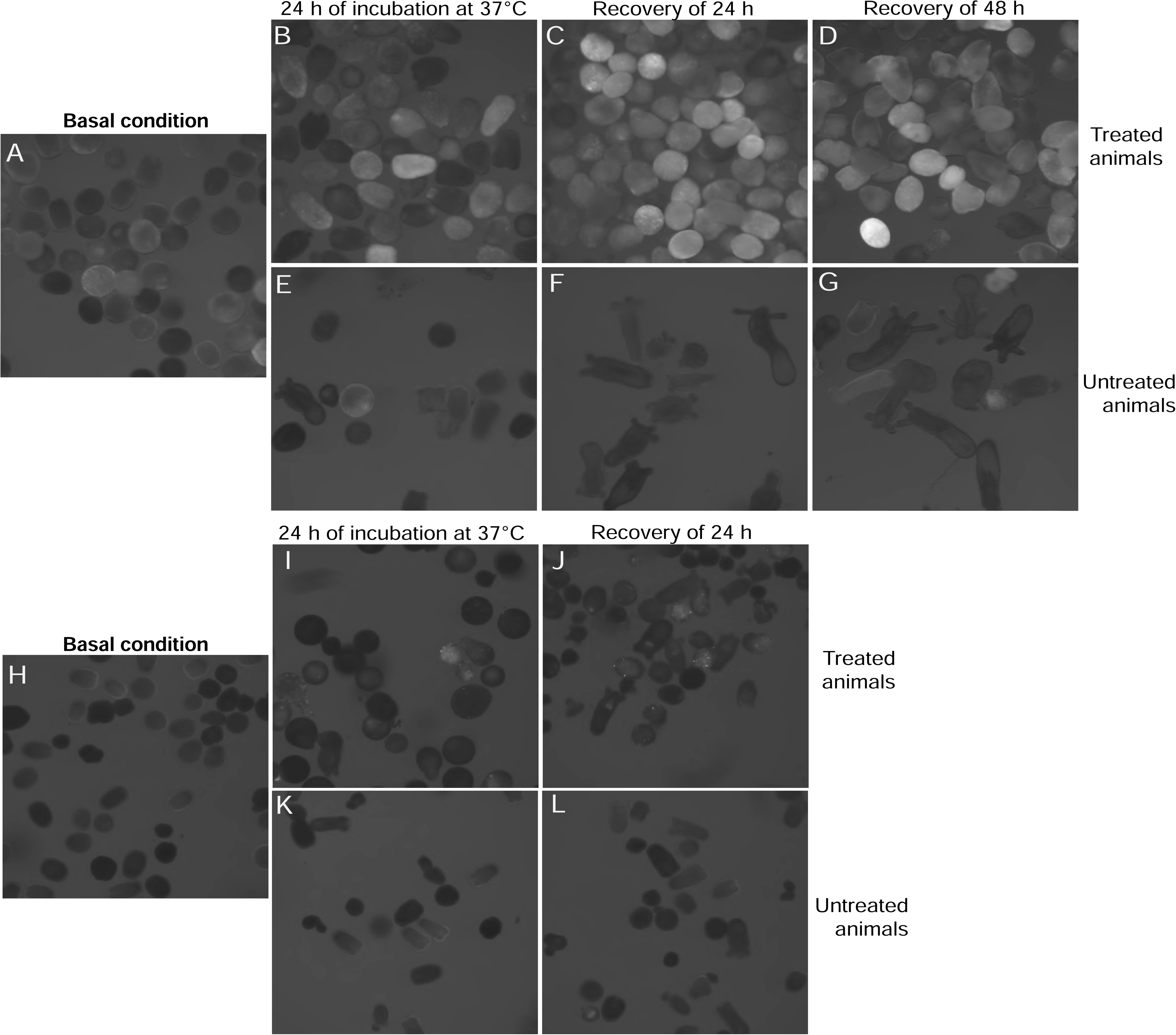
mCherry is upregulated under heat stress in the *RLRa* and *RLRb* transgenic reporter lines. 4-5 days post-fertilization *N. vectensis* larvae of the reporter lines were incubated for 24 hours at 37 °C. The larvae were given a recovery time of 19-48 hours at 22 °C. Untreated larvae from the same cross were used as a control group and were kept at 22 °C. (**A-G**) panels documenting the *RLRb* line and (**H-L**) panels documenting the *RLRa* line. (**A, H**) 4-5 days old larvae under basal conditions (22 °C) before the heat stress assay. (**B, I**) 5-6 days old larvae after 24 hours of incubation at 37 °C. (**C, D, J**) 6-7 days old larvae following recovery time of 19-48 hours at 22 °C after the incubation of 24 hours at 37 °C. (**E-G, K-L**) Untreated animals are used as control.

### Transcriptomic analyses highlight a link between stress and innate immunity that varies across populations

Intrigued by the sensitivity of the *RLR* promoters to stress, we explored previously obtained transcriptomic data from *N. vectensis* adult polyps that were incubated for 24 hours in harsh stress conditions which combined heat (37 °C) and salinity (40 ‰), as well as UV light for 6 hours (Sachkova, et al. 2020). Indeed, more than 20 genes were significantly upregulated under the stress conditions in both populations (**Fig. 3A-C**). Many of the upregulated genes were homologs of known mammalian Interferon Stimulated Genes (ISGs), including *OAS1*, *MAVS*, *RLRa*, *RLRb*, *IRF2-like3*, *GBP6-like*, *GBP3-like*, and *IRF4-like1*. These genes showed log_2_ fold changes ranging from 2 to 4 (Supporting Information tables S9 and S10). These findings align with our reanalysis of data from Weizman et al., 2021, where 21 genes were upregulated when comparing control and heat-stressed groups in the field-preconditioned population (Supporting Information table S11). In contrast, in the laboratory-preconditioned population, only 4 of the 56 immune marker genes (*OAS1*, *TRAF3-like, TRIM33-like1 1*, and *NFKB1)* were upregulated under heat stress (Supporting Information table S12). Notably, the anemones in this experiment were not from the common lab strain, which originates from Maryland, but from strains that originate from Massachusetts (both Sachkova et al. 2020 and Weizman et al. 2021) and North Carolina (Sachkova et al. 2020). Unexpectedly, despite the high similarity in the transcriptional response to stress between the two populations included in the Sachkova et al. (2020) study, transcription of several genes was altered under stress conditions in one population, but not in the other: for example, the homolog of *MCL1* (*myeloid cell leukemia 1*), a gene known to play an important role in cell homeostasis and bypass of apoptotic reaction to stress and infection, was upregulated in the North Carolina population but not in Massachusetts. An opposite trend, upregulation in the Massachusetts strain under stress and lack of response in the North Carolina strain, was noticed for the homolog of *IAP2* (*inhibitor of apoptosis 2*), a gene that in mammals affects the apoptotic response to infections and was shown to be stress-responsive (Dong, et al. 2002). These intriguing trends demonstrate that the immune response genes of different *Nematostella* populations reacts differently to the same stress conditions.

**Figure 3.**
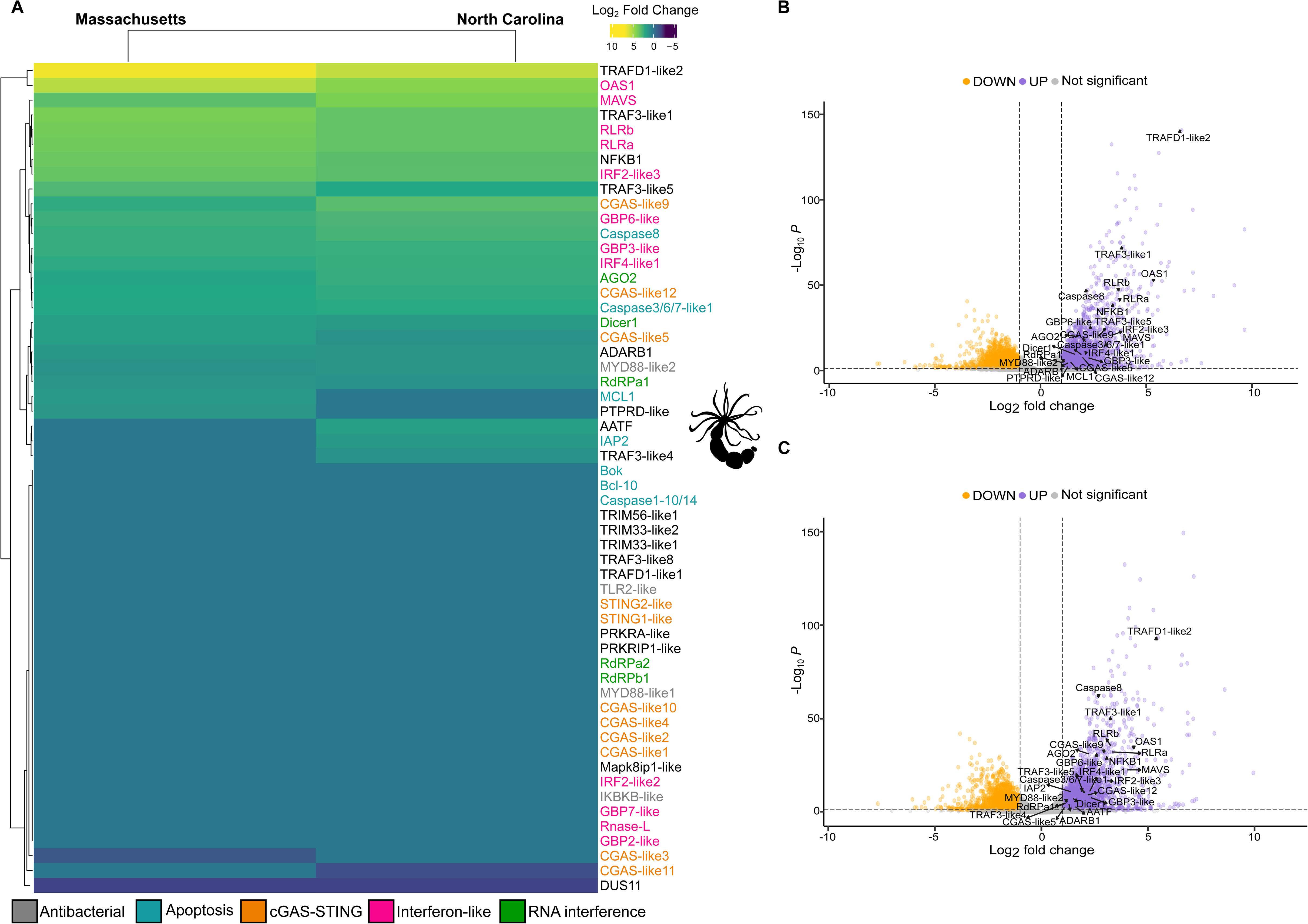
Transcriptional response of immune-related genes in *N. vectensis* under environmental stressors, including heat stress. (**A**) Heatmap showing the log_2_ fold change of immune-related genes in response to stress, comparing control and stress conditions in Massachusetts and North Carolina populations. These 56 immune genes were selected based on their known upregulation after poly(I:C) injection in *N. vectensis* zygotes (Lewandowska et al. 2021). The gene names were labeled with different colors representing the immune pathways they are involved in. (**B**) and (**C**) volcano plots showing differentially expressed genes (DEGs) in *N. vectensis* under combined environmental stress (Sachkova et al. 2020). Upregulated and downregulated genes (adjusted p-value < 0.05 and absolute log2 fold change > 1) are highlighted in purple and orange, respectively.

To test whether similar responses occur in other hexacorallians, we analyzed previously obtained transcriptomic data from experiments where the sea anemone *E. diaphana* and the stony coral *S. pistillata* were exposed to heat stress conditions (Cleves, et al. 2020; Savary, et al. 2021). Strikingly, in both the coral (**Fig. 4A-B**) and the sea anemone (**Fig. 4C-D),** immune-related genes exhibited significant differential expression. It was noticeable that unlike in *E. diaphana* and *N. vectensis*, in *S. pistillata* a portion of the responsive genes were downregulated under stress rather than upregulated (**Fig. 4A-B**). More specifically, 12 genes (*MAVS, RLRb, DCR1, RdRPa1, AGO2 STING2-like, Caspase8, TRAF3-like8, TRAF3-like1_1, TRAF3-like5_1, NFKB1* and *MYD88-like1*) were significantly upregulated when compared to a stress temperature of 34.5 °C (**Fig. 4A**, Supporting Information table S13), while 8 genes (*TRAF3-like8, TRAF3-like1_1, Caspase8, NFKB1, TRAF3-like5_1, TRAF3-like5_2, MAVS* and *STING2-like*) showed upregulation in comparison to a temperature of 32 °C (**Fig. 4A**, Supporting Information table S14). However, no genes were upregulated when comparing the control temperature to a stress temperature of 29 °C (**Fig. 4A**). Additionally, 5 genes (PRKRA-like, TRAF3-like5_2, RdRPb1, PRKRIP1-like and CGAS-like1) were downregulated when comparing the control temperature to 34.5 °C (**Fig. 4A**, Supporting Information table S13).

**Figure 4.**
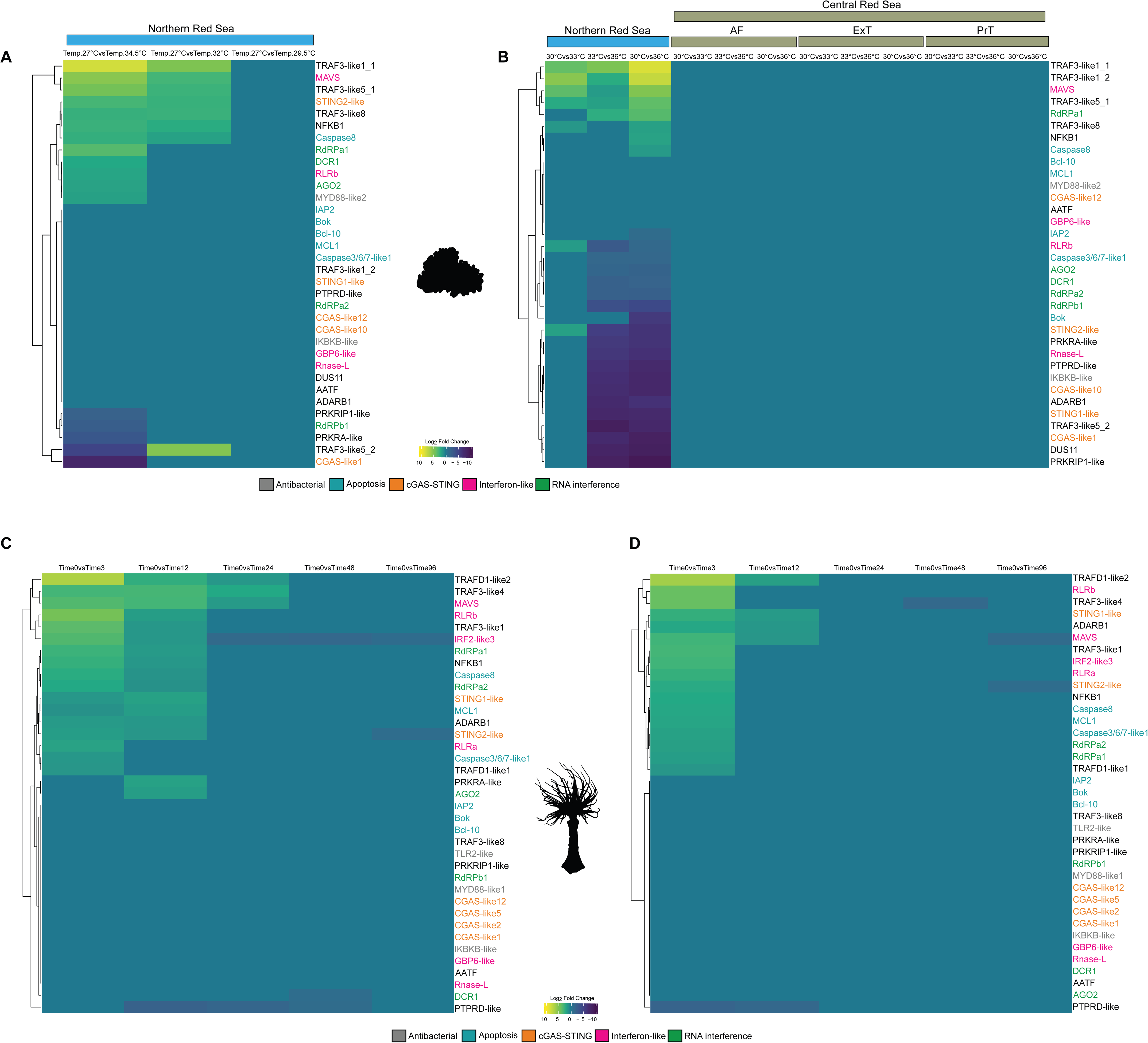
Immune response in *S. pistillata* and *E. diaphana* under heat stress. (**A**) and (**B**) Heatmaps showing the log_2_ fold change of immune-related genes in response to heat stress in S*. pistillata.* The data are derived from two previously published studies comparing baseline and stress temperatures in populations from the Northern and Central Red Sea. The populations in the Central Red Sea include Al Fahal Reef (AF), the ocean-facing exposed site of Tahala Reef (ExT), and the land-facing protected site of Tahala Reef (PrT). (**C**) and (**D**) heatmaps showing the log_2_ fold change of immune-related genes in response to heat stress, comparing baseline and stress temperatures across different time points from a previously published study on aposymbiotic (**C**) and symbiotic (**D**) *E. diaphana*. The gene names were labeled with different colors to represent the pathways related to the immune system. Statistical analysis using t-tests showed no significant differences (p < 0.05) between time 0 and time 3 conditions.

In the original experiment done on *E. diaphana*, Cleves et al. (2021) tested separately the transcriptomic response to heat stress of symbiotic (carrying zooxanthellae) and aposymbiotic (not carrying zooxanthellae) anemones. Anemones in this experiment were held at a control temperature of 27 °C and a stress temperature of 34 °C for different times. We identified upregulated genes only when comparing time 0 (control temperature) with 3 hours and 12 hours of exposure to the stress temperature in both aposymbiotic and symbiotic individuals (**Fig. 4C-D**, Supporting Information tables S18 and S19). Additionally, three genes remained upregulated after 24 hours of exposure in the aposymbiotic individuals, i.e, the response of these genes in the symbiotic group was much shorter lived than in the aposymbiotic group (**Fig. 4C-D**).

In another transcriptomic study (Voolstra, et al. 2021) that we reanalyzed the response to heat stress was documented separately for *S. pistillata* from the GoA vs. the same species from the CRS. Strikingly, for the 32 bona fide immune-response genes, in the CRS population we could find no significant differential expression, whereas in the GoA population we could find most of the genes being significantly upregulated or downregulated (**Fig. 4B, Supporting Information table S15-S17**), similarly to the trends reported for the study by Savary et al. (2021) that also focused on the GoA population (**Fig. 4A**).

### Characterizing the viral and antiviral transcription dynamics

In order to test whether the transcriptional immune response under stress is driven, at least partially, by an increase in viral load, we decided to first analyze existing data from *N. vectensis* and *E. diaphana*. As the stress experiments in *N. vectensis* were performed on strains different from the common lab strain where the core virome was already characterized (Lewandowska, et al. 2020), we started by assembling the viromes of the Massachusetts and North Carolina strains, following the same methodology we previously used for the common lab strain (Supporting Information tables S4 and S5). Then, we performed differential expression analysis for the assembled viral sequences. While in both populations there was a noticeable decrease in viral load under stress conditions, it was not significant (Supporting Information **Fig. 1A-B, Supporting Information table S20**).

We used the previously assembled viral sequences of *E. diaphana* (Bruwer and Voolstra 2018) and existing transcriptomic data from a heat stress experiment (Cleves, et al. 2020) to test whether viruses exhibit a difference in load under stress conditions in this species. However, we could not detect any significant trend in either symbiotic or aposymbiotic *E. diaphana* (Supporting Information **Fig. 1C-D**). We speculated that the failure in obtaining significant results may originate from the fact that we could map only very few reads to the assembled viruses. This in turn, might be due to many viruses not being polyA tailed as the sequencing methods used were utilizing polyA selection and/or due to the viral sequences being vastly outnumbered by host transcripts.

To circumvent these potential limitations, we decided to use a targeted approach where we amplified by specific primers six viral sequences representing the most dominant members of the core virome of *N. vectensis* and six major antiviral genes (*RLRa*, *RLRb*, *IRF2-like3*, *OAS*, *DCR1* and *AGO2*) by quantitative PCR (qPCR; Supporting Information table S2). This approach also allowed us to assay the dynamics of the stress response. We incubated adult *N. vectensis* polyps of the common lab strain to heat stress of 37 °C and with samples taken at several time points. After RNA extraction and cDNA synthesis, we used qPCR to measure viral sequences as well as host antiviral gene expression. Unexpectedly, four of the viruses showed a significant load increase within 30 minutes from the beginning of the incubation and five of them peaked at 3 hours **(Fig. 5A)**. The antiviral genes exhibited mild, yet significant, upregulation only after 1-3 hours, and most of them peaked at 24 hours after the start of the incubation under stress conditions (**Fig. 5B**). Interestingly, variation in the dynamics of the responses was observed both between the antiviral genes and between the viral sequences (**Fig. 5A-B**).

**Figure 5.**
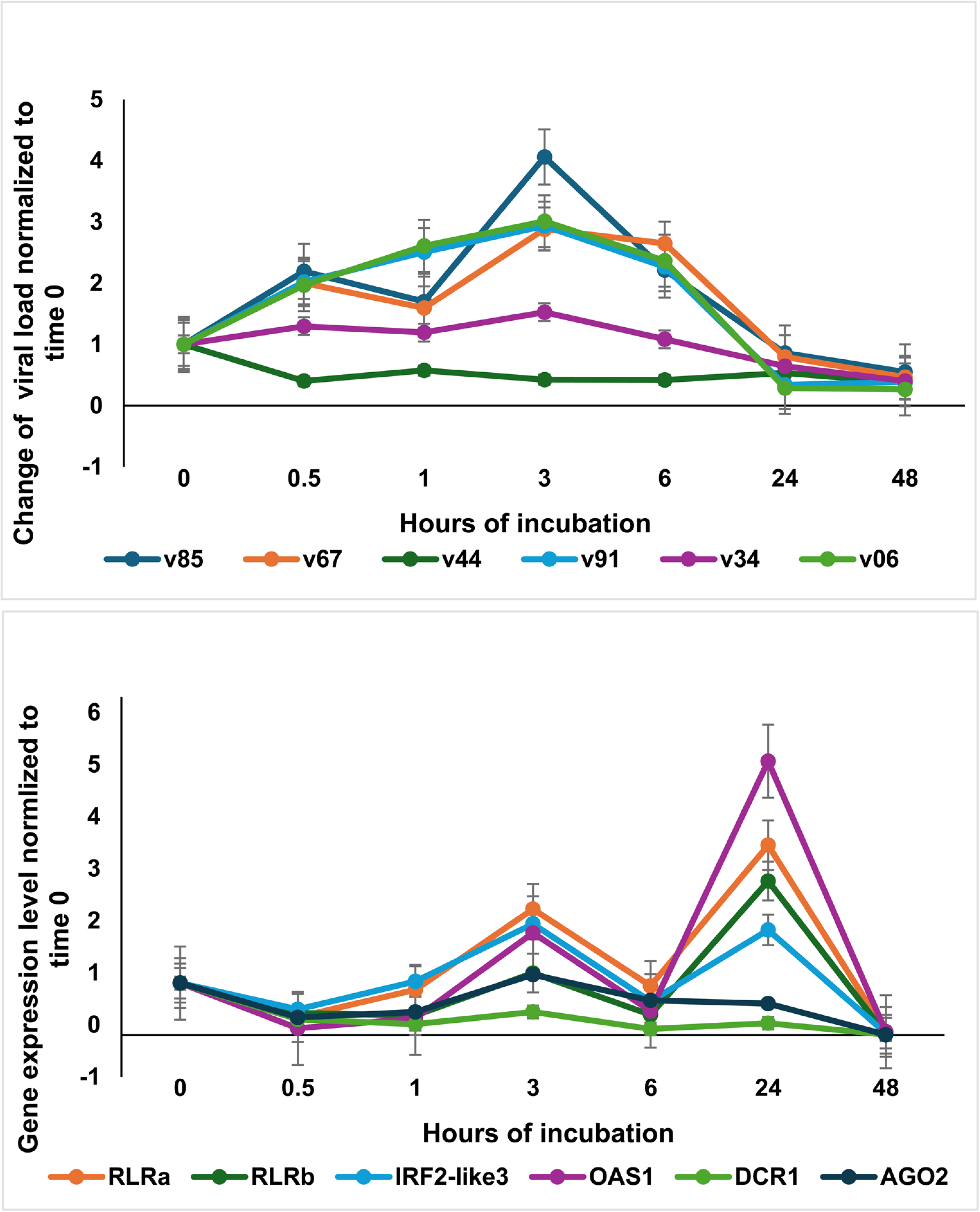
Dynamic heat stress assay for testing the antiviral immune response compared to the change of the viral load in *N. vectensis*. Adult polyps were incubated at 37 °C and assayed at different time points for the expression levels of six antiviral genes (**A**) (Lewandowska et al. 2021) compared to six sequences from the *N. vectensis* core virome (**B**) (Lewandowska et al. 2020). The changes were measured by qPCR (see methods).

## Discussion

In this work, we have set to explore how stress and immune response are linked in hexacorallians. In addition to demonstrating this link at the transcriptional level in two sea anemone species and one species of stony corals, we unexpectedly revealed that the symbiotic state in *E. diaphana*, as well as the geographic origin in *N. vectensis* and *S. pistillata*, affect the transcriptional response (**Fig. 3 and Fig. 4**). We already reported transcriptional and proteomic differences in the response to stress between the northern and southern *N. vectensis* populations in the context of venom production (Sachkova, et al. 2020). This suggests that these populations have transcriptionally adapted to different environmental conditions that typify their habitats, which is plausible in light of the extreme differences in the average temperatures and sunlight exposure between those habitats. For example, between March-December 2016, in 114 days the water temperature at the North Carolina habitat crossed 36 °C whereas this happened in Massachusetts only in 39 days during the same period (Sachkova, et al. 2020).

Another notable result of our analysis is that the set of immune-related genes in *S. pistillata* from the CRS is unresponsive to heat stress while the same set responds in the population of the same coral species from the GoA (**Fig. 4A-B**), in line with previous findings that CRS corals exhibit a measured response. Interestingly, CRS corals showed a consecutive upregulation of certain stress genes also known as frontloading (Voolstra et al 2021). Corals of the Red Sea are exceptionally heatDresistant, yet bleaching events have increased in frequency in the central and southern sections (Monroe, et al. 2018; Genevier, et al. 2019). By comparison, thus far, the northern Red Sea remains largely “bleaching-free” and is sometimes presented as a “coral sanctuary” during climate change (Fine, et al. 2013; Osman, et al. 2018).

Our transcriptomic analysis suggests that at least a partial explanation for this difference between the GoA and CRS populations may be their response to heat stress by modulating the immune system. This may be a crucial component in retaining zooxanthellae under harsh climatic conditions, especially as downregulation of immune response was previously linked to hexacorallian symbiosis with these algae (Mansfield and Gilmore 2019). It is also relevant for the difference we found between the transcriptomic responses to heat stress of immune-related genes in aposymbiotic (**Fig. 4C**) and symbiotic (**Fig. 4D**) *E. pallida*, where the former group displays a prolonged response.

For discerning between the two alternative scenarios: (i) heat stress induces viruses that in turn induces the immune response or (ii) the immune system of hexacorallians evolved to be upregulated upon heat stress, posited that this would be advantageous against viruses that are activated by heat stress, we measured selected viruses (**Fig. 5A**) and antiviral response (**Fig. 5B**). As we detected increase in the viral load prior to the host antiviral response in *N. vectensis*, it is plausible that the extremely swift response of the viruses to the heat stress contributes to promoting the host immune response that is lagging behind. This rapid viral activation and the consequential raid antiviral innate immunity response by the host also make it unlikely that viral loading is a secondary response of enduring stress.

Altogether, our study reveals that diverse hexacorallians separated from one another by hundreds of millions of years of evolution respond to heat stress by modulating their immune system in a matter of mere hours and that viruses putatively play a role in this response by responding even faster to the stress. Future work could focus on further untangling the effects of the stress on the host immune system from the response of the viruses, characterize the potential link to bleaching, and hopefully provide deeper understanding of the local adaptation of hexacorallian populations and its ultimate underlying causes.

## Acknowledgements

The authors acknowledge the help provided by the personnel of the Genomics Technologies Center of the Alexander Silberman Life Sciences Institute of the Hebrew University. This work was supported by the European Council Research consolidator grant AntiViralEvo 863809 to Y.M.

## Conflict of interest statement

The authors do not have any conflicts of interest to declare.

## Data availability statement

All the data used for this work is available either in the main text or in the Supporting Information files. All sequence data are derived from public databases mentioned in the text.

**Supporting Information Figure 1. Differences between viral reads in *N. vectensis* and *E. diaphana* under stress.** While *N. vectensis* polyps from both Massachusetts (**A**) and North Carolina (**B**) exhibited a decrease in reads mapping to viral sequences under heat stress conditions, the difference was not statistically significant. In *E. diaphana* no significant difference was detected in the number of viral reads in both aposymbiotic (**C**) and Symbiotic (**D**) polyps when anemones kept for 3 hours under heat stress conditions (Time 3) were compared to those kept under control conditions (Time 0).

**File1. Virus sequences from *N. vectensis* core virome (Lab population Maryland) were traced for testing the increase in the viral load.**

**File2. Virus sequences from *N. vectensis* virome (populations of North Carolina and Massachusetts) and core virome were traced for testing the increase in the viral load.**

**Table S1. NveRLRa and NveRLRb promotor region primers, that are used for cloning and the generation of the reporter transgenic lines with mCherry.**

**Table S2. The sequence of primers used for qPCR.**

**Table S3. Statistical t-test for the dynamic heat stress assay. *N. vectensis*.**

**Table S4. List of orthologs identified for *E. diaphana* and *S. pistillata* through orthology analysis with OrthoFinder, based on homologs established in Nematostella vectensis.**

**Table S5. Protein sequences of orthologous immune system-related genes were obtained for the three species analyzed in this study (*N. vectensis*, *E. diaphana*, and *S. pistillata*).**

**Table S6. Core virome sequences from *E. diaphana*.**

**Table S7. Read counts obtained from mapping against the virome sequences from *N. vectensis*. (Massachusetts and North Carolina).**

**Table S8. Raw read counts obtained from mapping against the core virome sequences from *E. diaphana*. (Aposymbiotic_Time0vsTime3 and Symbiotic_Time0vsTime3).**

**Table S9. Differential expression of immune-related genes in *N. vectensis* (Massachusetts population).**

**Table S10. Differential expression of immune-related genes in *N. vectensis* (North Carolina population).**

**Table S11. Differential expression of immune-related genes in *N. vectensis* (Field population).**

**Table S12. Differential expression of immune-related genes in *N. vectensis* (Laboratory population).**

**Table S13. Differential expression of immune-related genes in *S. pistillata* (27°C vs 34.5°C).**

**Table S14. Differential expression of immune-related genes in *S. pistillata* (27°C vs 32°C).**

**Table S15: Differential expression of immune-related genes in *S. pistillata* (30°Cvs 33°C).**

**Table S16: Differential expression of immune-related genes in *S. pistillata* (30°Cvs 36°C).**

**Table S17. Differential expression of immune-related genes in *S. pistillata* (33°Cvs 36°C).**

**Table S18. Differential expression immune-related genes in *E. diaphana* (Aposymbiotic) (Time 0h versus Time 3h and Time 0h versus Time 12h).**

**Table S19. Differential expression immune-related genes in *E. diaphana* (Symbiotic) (Time 0h versus Time 3h and Time 0h versus Time 12h).**

**Table S20. Statistical t-test. *N. vectensis* of Massachusetts and North Carolina Populations.**

## Notes

### Competing Interest Statement

The authors have declared no competing interest.

