## Supplementary Figure 1 for "Heat stress drives rapid viral and antiviral innate immunity activation in Hexacorallia"

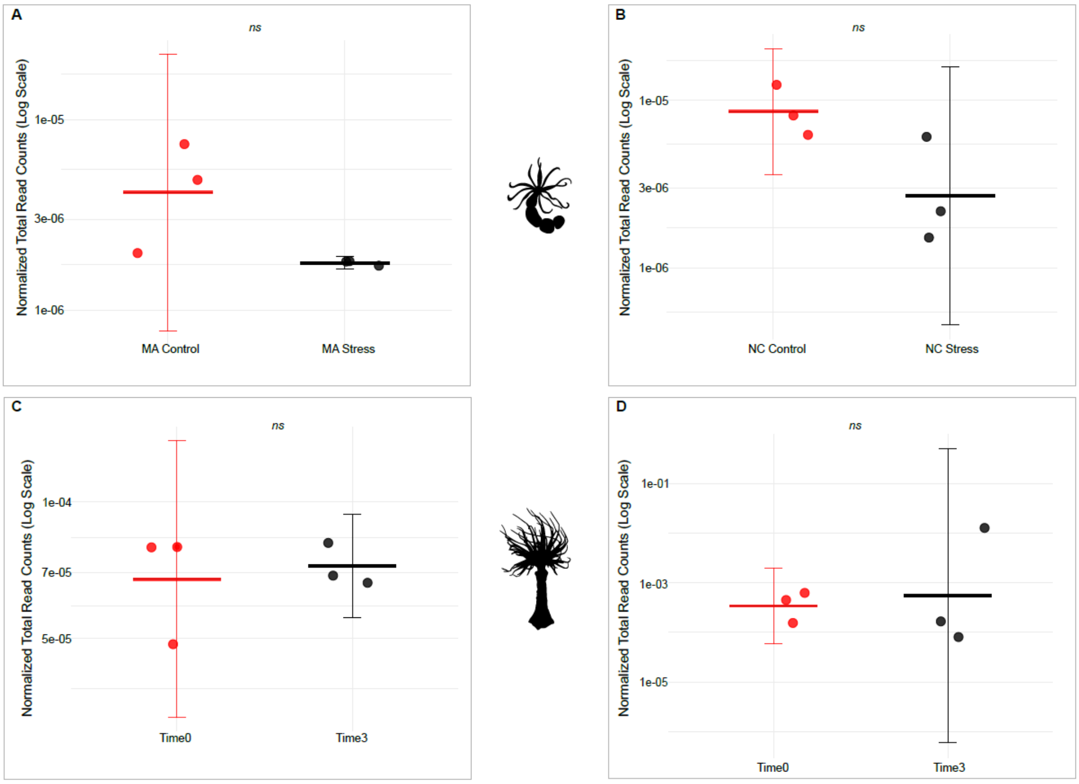


**Figure S1. Viral read count analysis in *Nematostella vectensis* and *Exaiptasia diaphana* under environmental stressors, including heat stress.**

(A,B) Total normalized viral read counts in control and stress conditions for Massachusetts (A) and North Carolina (B) populations. (C, D) Total normalized viral read counts under Time0 and Time3 conditions for aposymbiotic (C) and symbiotic (D) *Exaiptasia diaphana*.Statistical analysis using t-tests showed no significant differences ($p > 0.05$) between control and stress conditions within each population. Error bars indicate the standard error. *ns*=non-significant.
